# *Monkey See, Model Knew*: Large Language Models Accurately Predict Visual Brain Responses in Humans *and* Non-Human Primates

**DOI:** 10.1101/2025.03.05.641284

**Authors:** Colin Conwell, Emalie McMahon, Akshay Jagadeesh, Kasper Vinken, Saloni Sharma, Jacob S. Prince, George A. Alvarez, Talia Konkle, Margaret Livingstone, Leyla Isik

**Affiliations:** Johns Hopkins University, Department of Cognitive Science; Harvard Medical School, Department of Neurobiology; Harvard University, Department of Psychology

## Abstract

Recent progress in multimodal AI and ‘language-aligned’ visual representation learning has rekindled debates about the role of language in shaping the human visual system. In particular, the emergent ability of ‘language-aligned’ vision models (e.g. CLIP) – and even pure language models (e.g. BERT) – to predict image-evoked brain activity has led some to suggest that human visual cortex itself may be ‘language-aligned’ in comparable ways. But what would we make of this claim if the same procedures could model visual activity in a species without language? Here, we conducted controlled comparisons of pure-vision, pure-language, and multimodal vision-language models in their prediction of human (N=4) and rhesus macaque (N=6, 5:IT, 1:V1) ventral visual activity to the same set of 1000 captioned natural images (the ‘NSD1000’). The results revealed markedly similar patterns in model predictivity of early and late ventral visual cortex across both species. This suggests that language model predictivity of the human visual system is not necessarily due to the evolution or learning of language *perse*, but rather to the statistical structure of the visual world that is reflected in natural language.

## 1 Introduction

The idea that language shapes how we ‘see’ the world has long been one of the most actively debated ideas in cognitive (neuro)science (1; 2; 3; 4; 5), and has evolved through many forms over that time. A recent evolution of this idea has manifested in the form of competing hypotheses about the extent to which high-level human visual cortex is ‘language-aligned’ – or, in other words, the extent to which lingustic or linguistically-learned structure is evident in visual brain responses (6; 7). The resurgence of this debate is predicated in large part on two seminal findings in research on modern deep learning models: first, the finding that ‘language-aligned’ machine vision models (e.g. CLIP) are some of the most predictive models to date of image-evoked activity in the visual brain (8; 9); and second, the finding that even pure-language models (e.g. BERT) are capable of predicting image-evoked brain activity by way of image captions alone (10; 11).

Here, we apply a logical razor to this debate in the form of assessing whether these two key findings hold in the brain of a species that does not have language. We call this the ‘monkey razor’, and define it as follows: If the ability of ‘language-aligned’ vision models or pure-language models to predict image-evoked brain activity is indeed evidence of language having (re-)shaped visual representation, we should not find similar predictivity in monkeys.

We used encoding models fit to the feature spaces of a diverse set of pure-vision, pure-language (LLMs), and multimodal (language-aligned) vision (VLMs) models to predict image-evoked brain activity in the ventral stream of 4 humans and 6 rhesus macaques shown the same set of 1000 natural images from the Natural Scenes Dataset (NSD) (12). The brain-likeness of the pure-vision and language-aligned vision models was assessed on the images themselves. The brain-likeness of the pure-language models was assessed using an average of the embeddings for the first 5 captions associated with each image (collected from metadata for the MS-COCO dataset (13), from which NSD images are curated).

We find that vision and language models provided accurate predictions of neural responses in highlevel ventral stream regions of both human and macaque visual cortex. The majority of variance explained by pure-vision and pure-language models in human OTC was shared with macaque IT. The remaining ‘uniquely human’ variance was equally well explained by the two model modalities. Together, these results suggest that language model predictivity of human visual cortex is likely not due to language learning or true representations of language, but instead a reflection of a convergence between vision and language models based on the large-scale, end-to-end statistical learning that defines them both.

## 2 Results

### General Approach

The encoding models for the human (fMRI) brain activity were fit to reliability-selected voxels (*NCSNR >* 0.2) in a broad mask of early visual cortex (EVC, N=15326 voxels) and occcipitemporal cortex (OTC, N=29840 voxels), with both anatomical and functional criteria as the basis of inclusion. The encoding models for the monkey (electrophysiology) brain activity were fit to multi-unit responses (i.e. average firing rates in a 150ms window) from arrays placed either in macaque V1 (N=34 units) or inferotemporal (IT) cortex (N=394 units). Details on all aspects of our approach are available in Section 4.

### Vision and Language Models Predict High-Level Regions in Human Ventral Stream

Commensurate with previous findings (14; 10), we found that pure-language model embeddings over image captions could predict high-level human ventral visual activity almost as accurately as pure-vision models (Figure 1B: Top Row, Table 1: Row 1). There was not a substantial difference between pure-vision and language-aligned vision models in predicting OTC responses. In contrast, language models performed far worse than both types of vision models in predicting early visual cortex activity (Table 1: Row 3). The poor performance of these models in early visual cortex provides a sanity check, demonstrating that the LLMs are not just statistically overpowered in general.

**Table 1:**
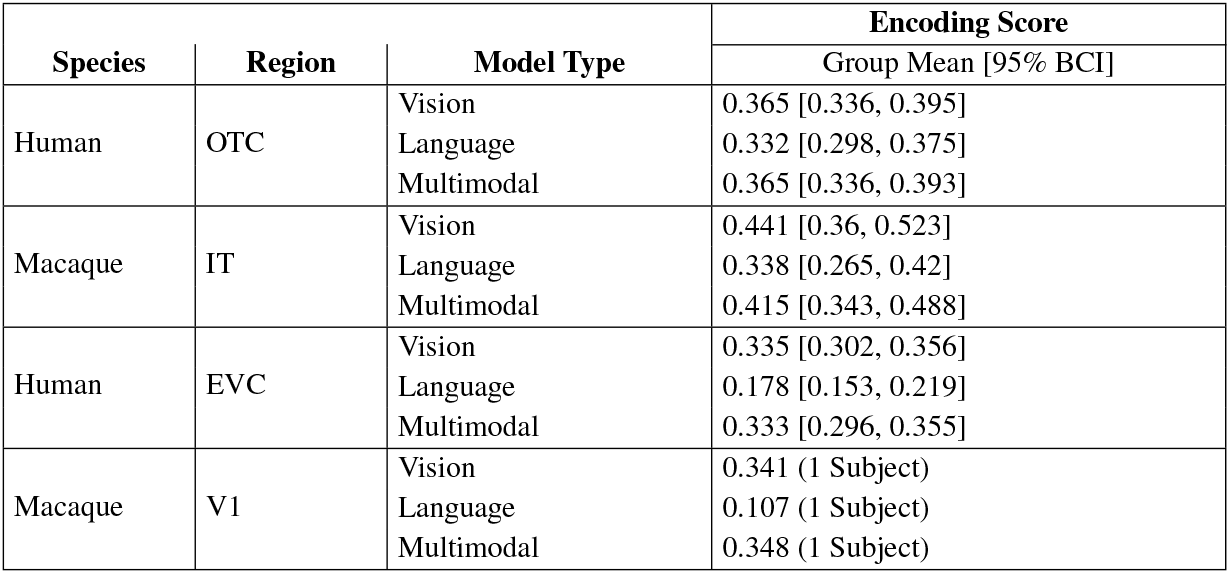
Human-Monkey-Model Comparisons Summarized

**Figure 1:**
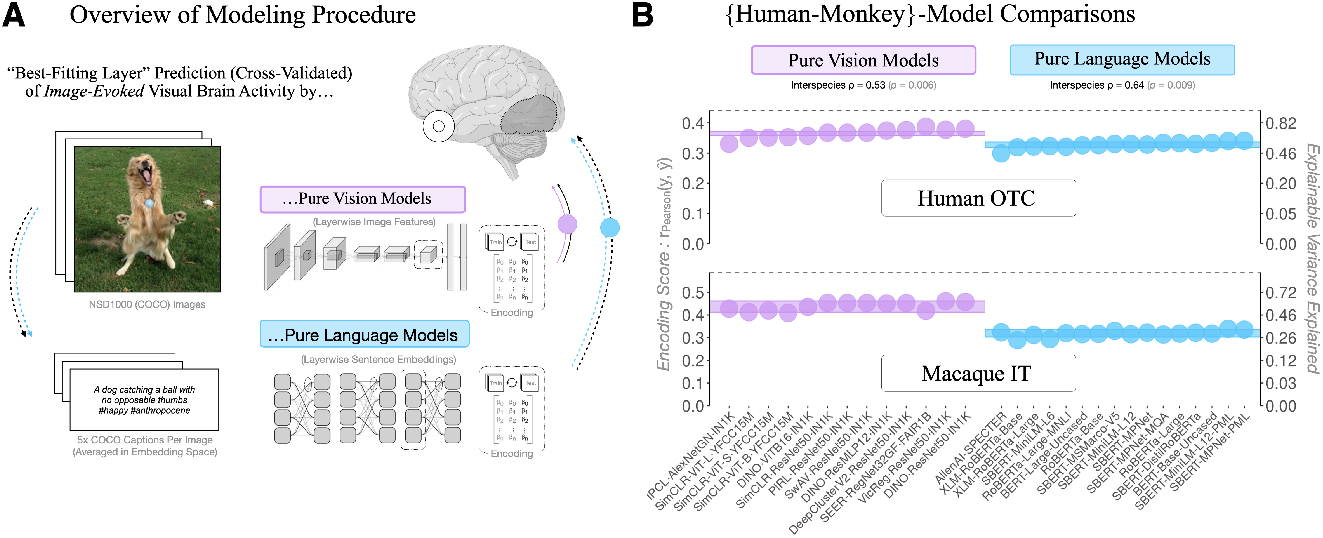
(A) Overview of modeling procedure: *Image-evoked visual brain activity* was predicted from the image (for vision models) or the image captions (for language models), using a unit-wise (voxel / neuron) encoding analysis. (B) Encoding accuracies from the most brain-like layer of a series of (unimodal) pure vision and pure language models in prediction of both human occipitotemporal cortex (OTC) and macaque inferotemporal cortex (IT). Individual points are the accuracies for each model. The horizontal, semitranslucent rectangles extending across the dots are the means *±*95% bootstrapped CIs (across models) per model type (modality).

### Vision and Language Models Predict High-Level Regions in Macaque Ventral Stream

Applying the same encoding procedures with the same stimuli to prediction of brain activity in macaque visual cortex, we found that, as in humans, pure-language models are remarkably accurate in predicting high-level ventral visual activity (Figure 1B: Bottom Row, Table 1: Row 2). Also as in humans, we found that pure-language models perform poorly in prediction of early visual cortex (Table 1: Row 4). There was also no substantial difference between pure-vision models and language-aligned vision models. There was, however, a slightly more pronounced advantaged of pure-vision over pure-language models in macaque IT (0.441 versus 0.338) compared to human OTC (0.365 versus 0.332). A time-resolved encoding analysis suggests that the pure-vision and pure-language models have a similar time course of predictivity, further underscoring the overlap between the learned representations across modalities (Supplementary Figure A1).

### Better Models of Human Ventral Stream are Also Better Models of Macaque Ventral Stream

The results above demonstrated that vision and language models perform largely similarly across the species; but what about the more granular comparison instantiated by the *rank-order* correlation of models across the species (Figure 1B). For pure-vision models, this correlation was ρ = 0.53 [0.49, 0.56] (*p* = 0.006), suggesting individual vision models are similarly predictive of both species. For pure-language models, this correlation was 0.64 [0.61, 0.65] (*p* = 0.009) – slightly, but significantly higher than the interspecies correlation for the vision models (with bootstrapped mean differences (Δρ) = 0.053 [.032, .076]; and differences *>* 0 in 976 / 1000 bootstraps, *p* = 0.023). In short, better language models of human visual cortex are also better language models of macaque visual cortex.

Accordingly, hypotheses about why some language models do better than others in their prediction of human visual cortex should apply commensurately to macaque visual cortex, as well.

### Uniquely Human Variance is Equally Well Predicted by Vision and Language Models

The slight difference in language versus vision models in predicting macaque IT relative to human OTC may be due to multiple factors, including the species-specific recording modalities and preprocessing steps (e.g., electrophysiology versus functional MRI) or differences in visual experience of the human versus macaque subjects.

The question of most relevance, here, though, is whether the difference is attributable primarily to language (or at least, language as encoded in the pure-language models). To assess this directly, we performeded a 3-way unique variance analysis, predicting human OTC activity with all seven combinations of the groupings of three predictors: macaque brain activity, vision and language model embeddings (using the model embeddings for the most OTC-predictive model from each set).

The goal of this analysis was to understand how well different types of models predict ‘uniquely human’ neural signals in OTC. The logic being that if the difference between humans and monkeys is a difference attributable to language (over and above vision), then the unique variance explained by language models should be greater than the unique variance explained by vision models. We find this was not the case (Figure 2, and Table 2). The majority (8.3% + 12.1% + 46.6% = 67%) of the explainable variance in human OTC was shared with macaque IT and at least one of the models. We note there was almost no shared variance between the macaque and human data that was not also shared with one of the models. Of the remaining ‘uniquely human’ variance, a similar amount was attributable to the vision (18.4%) and language language (14.6%) models, with slightly more unique variance explained by the vision model. The relative symmetry of the unique variances suggests that the difference between human OTC and macaque IT is not a function of language, but may instead be a reflection of any number of the modes of difference in species, recording modality, or experimental setup.

**Table 2:** Interspecies Unique Variance Analysis Summarized

| Regression | Unique |  |  | Shared (2-Way) |  |  | Shared (3-Way) |
| --- | --- | --- | --- | --- | --- | --- | --- |
|  | Species | Vision | Language | Species+ Vision | Species+ Language | Vision+ Language | All Predictors |
| Human ∼ Macaque | $\sim.0\%$ | 18.4% | 14.6% | 8.3% | 12.1% | $\sim.0\%$ | 46.6% |
| Macaque ∼ Human | $\sim.0\%$ | 11.8% | $\sim.0\%$ | 11.5% | 25.4% | 13.1% | 38.2% |

**Figure 2:**
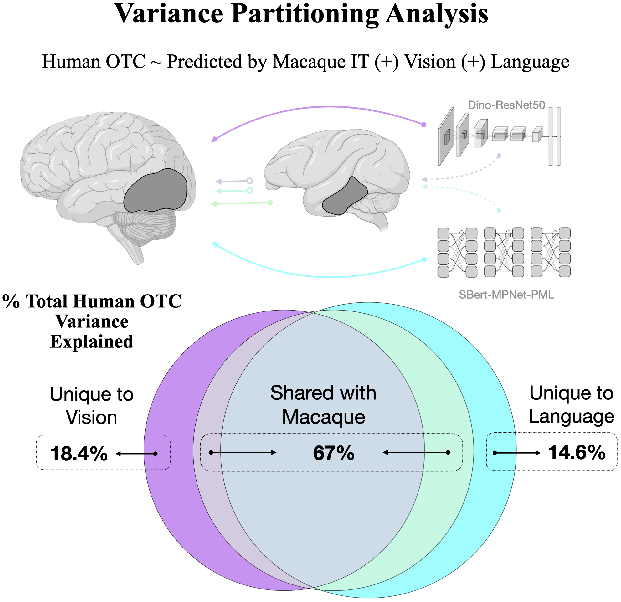
Results from a 3-way variance partitioning analysis designed to quantify the variance in human visual brain activity attributable uniquely to vision and uniquely to language, all while accounting for the variance shared with macaque visual brain activity. This figure shows the variance partitioning of human OTC across the 3 key predictors for this analysis: macaque IT, the most-predictive pure vision-model (Dino-ResNet50) and the most predictive pure-language model (SBERT-MPNet-PML). Most of the variance in this analysis is shared across both macaques and models; the variance attributable uniquely to either or the models is comparable across vision and language – with a slight advantage to vision as the greater source of ‘uniqueness’ between the species.

Voxel-wise maps of the variance unique to humans and each model modality reveal slightly different anatomical patterns however, with vision models explaining more variance unique to humans in posterior regions, and language models in anterior and lateral regions (Figures A3-A6).

## 3 Discussion

Here we show that the ability of ‘language-aligned’ vision models and pure-language models to predict image-evoked brain activity in human high-level visual cortex is likely not evidence of language having reshaped vision. We find similar trends using these models to predict visual brain responses in non-human primates, and demonstrate that those differences which do exist between humans and non-human primates are not directly attributable to the structures of language, as captured by LLM embeddings, alone. Such is the nature of the ‘monkey razor’: If, *caeteris paribus*, an experimental effect holds in both humans and monkeys, that effect cannot be attributable to the structure, function, deployment, or learning of language *per se*. Thus, the more likely explanation is that the representational overlap between pure-language models and the high-level primate visual brain reflects a structure learnable in large part through the hierarchical encoding of natural image statistics. Language (as learned by language models) may approximate the representational endpoints of this process, but only to the extent that these statistics are reflected in the language we use to describe the world around us (a world the language models themselves cannot actually ‘see’).

The observations we have made in primate brain prediction align well with recent findings in AI research that show vision and language models seem to be converging on common – ‘platonic’ (15) – representations (16; 17), even in the absence of explicit alignment. And while we note that this particular alignment (between vision and language models) may be more a function of the effective overlap in the training data of the internet than of shared world knowledge, it does seem to suggest again that language reflects visual structure. However, recent research suggests that this shared structure between vision and language is limited to pictoral visual descriptions of a scene (18), and that language models predictivity of visual cortex does *not* extend to linguistic descriptions of dynamic events (19) or more abstract conceptual features of an image (20).

Interestingly, we do find a small but consistent difference in the voxel-wise patterns of “uniquely human” variance explained by vision versus language models (Figures A3-A6). Of note, language models are slightly more predictive in more anterior and lateral vision regions, particularly the extrastriate body area (EBA), consistent with prior research showing an advantage of language-aligned vision models in these regions (8). The EBA is involved in social processing and has been recently shown to represent configural information about bodies and objects in a scene (21; 22). Such relational representations may explain the advantage of language models in this region, and these results suggest that perhaps some aspects of these representations may be unique to humans.

Further work is needed to make sense of the lingering difference, however small, between language model predictivity of human OTC and macaque IT. One major factor that merits further scrutiny here is the translation between different neural recording modalities: fMRI signals, for example, may include later visual components (including feedback) not evident in the electrophysiological signals. Differences in experimental setup and task demands (i.e. stimulus duration, ISI, and freeviewing versus fixation) may also affect the extent to which semantic content is captured in visual cortex (23). It is also important to note the drastically different visual diets of the human and macaque subjects, the latter having limited exposure to varied natural scene statistics. Finally, and perhaps most importantly, we should aspire to continue collecting visual brain data in both species that pushes the limits of representation learnable through image statistics alone—and extends more explicitly into the kinds of conceptual territories where the structures of language are most indispensable for understanding.

## 4 Materials and Methods

### 4.1 Human data

Human fMRI data were used from the Natural Scenes Dataset (12), which contains measurements of over 70,000 unique stimuli from the Microsoft Common Objects in Context (COCO) dataset120 at high resolution (7T field strength, 1.6-s TR, 1.8mm3 voxel size). We used the brain responses to the “NSD1000”, the set of 1000 COCO stimuli that overlapped between subjects, and limited analyses to the 4 subjects (subjects 01, 02, 05, 07) that saw these images with at least 3 repetitions. The 3 image repetitions were averaged to yield the final voxel-level response values in response to each stimulus. Beta responses were estimated using the GLMsingle toolbox (24), which implements optimized denoising and regularization procedures to accurately measure changes in brain activity evoked by experimental stimuli.

We performed voxel selection to improve signal-to-noise ratio (SNR) in our target data using a reliability-based voxel selection procedure (25) to select voxels containing stable structure in their responses. Specifically, we use the NCSNR (“noise ceiling signal-to-noise ratio”) metric computed for each voxel as part of the NSD metadata as our reliability metric. In this analysis, we include only those voxels with NCSNR *>* 0.2. After filtering voxels based on their NCSNR, we then filtered voxels based on region-of-interest (ROI) as described in prior work (26).

### 4.2 Macaque electrophysiology data

Electrophysiological multiunit activity was recorded from six macaque monkeys (5 macaca mulatta, 1 macaca nemestrina), using chronically implanted microelectrode arrays. In one subject, a 32-channel floating microelectrode array was implanted in primary visual cortex (V1). Three subjects were implanted with microwire brush arrays targeting anterior IT (aIT) cortex (64 channels, 64 channnels, 128 channels), and two subjects were implanted with microwire brush arrays (both 64 channels) targeting central IT (cIT) cortex. Cortical areas were localized using a combination of anatomical landmarks, structural MRI, and functional MRI. To consider the most analogous regions to human fMRI data, we grouped all IT recordings and report the average scores in V1 and IT.

Images were presented at the center of each subject’s population receptive field while subjects passively fixated on a point at the center of the screen. Subjects were rewarded for maintaining gaze at the fixation point for the duration of the image presentation. Images subtended 6 degrees of visual angle and were presented for 150ms with an interstimulus interval of 167 ms. Receptive fields were mapped in a prior session by rapidly presenting a separate set of small images (2 x 2 degrees size) at a dense grid of locations spaced apart by 1 degree. We presented a total of 1110 unique images to each monkey, including 110 fLoc functional localizer images (27) and 1000 COCO images drawn from the Natural Scenes Dataset (12). Each image was viewed a median of 19-40 times over six recording sessions by each subject to allow for trial-averaging responses across repetitions. We modeled the multi unit activity in each electrode in the 0-150 ms window following stimulus onset. For two monkeys (M1 and M2) we also modeled the time-resolved IT multi unit activity from 0-500 ms post stimulus onset in overlapping 150 ms bins, sampled every 10ms.

### 4.3 Model Selection

We tested vision-only, language-only, and vision-language deep neural networks.

Vision-only models included N=13 purely self-supervised visual contrastive-learning models from the VISSL model zoo (28), Dino-V1 variants (29), and one instance-level prototype contrastive learning (IPCL) model (30). Vision-language models consists of N=10 models from the OpenAI CLIP, OpenCLIP, SLIP, and PyTorch-Image-Models repository (31; 32; 33; 34). Our sample of large language models consisted of N=16 models from Hugging Face and the SBERT repositories (35; 36).

### 4.4 Model-brain mapping

#### 4.4.1 Feature extraction

We utilized DeepJuice (26), a python package in alpha-release that allows for memory efficient feature extraction from each layer of a DNN. We extracted the intermediate representations from every unique computational submodule (referred to here as layers) of every model.

We then used GPU-optimized sparse random projection (SRP) implemented in DeepJuice to project the activations in a 5920-dimensional feature space based on the Johnson–Lindenstrauss lemma with ε = 0.1. The sparse random projection matrix consists of zeros and sparse ones, forming nearly orthogonal dimensions, which are then normalized by the density of the matrix (the inverse square root of the total number of features). The layerwise feature maps are then projected onto this matrix by taking the dot product between them. The output of the procedure is a reduced layerwise feature space of size of 1000 images × 5920 dimensions with a preserved representational geometry. Note that in cases where the number of features is less than the number of projections suggested by the JL lemma, the original feature map is effectively upsampled through the random projection matrix, again yielding a matrix of 1000 × 5920 dimensions.

#### 4.4.2 Unit-WISE LINEAR ENCODING

We computed a linear encoding model for each unit (voxel or electrode) as a weighted combination of the 5920 model dimensions, using brain data from our training set of 500 images. The DeepJuice fitting procedure for each voxel leverages a custom GPU-accelerated variant of SciKit-Learn’s cross-validated ridge regression function (‘RidgeCV’), a hyperefficient regression method that uses generalized cross-validation to provide a LOOCV prediction per image (per output).

This fit was computed over a logarithmic range of alpha penalty parameters (1e-1 to 1e7), to identify each unit’s optimal alpha parameter. The RidgeCV function is modified in DeepJuice in order to select the best alpha using Pearson correlation as a score function (the same score function used for overall model evaluation), and to parallelize an internal for-loop for greater efficiency. This yielded a set of encoding weights for each unit.

We use these weights to predict unit-wise responses in the held-out 500 test videos. Encoding score is calculated by correlating actual and predicted unit-wise responses. Unit-wise scores are then averaged in each region (e.g., human OTC or macaque IT).

### 4.5 Variance Partitioning

To test how much of the difference between vision and language models in the prediction of macaque visual cortex is a difference attributable *uniquely* to language, we performed a 3-way variance partitioning analysis, closely following a methodology outlined by Lescroart et al. (37) and tutorialized by Tarhan (38). This 3-way analysis entails the fitting of 7 cross-validated regressions, always with one of the two species’ visual brain activity (human or macaque) as the primary outcome, but with different permutations of 3 predictors (the brain activity of the other species, the best-performing pure vision model, and the best-performing pure language model). Taking human brain activity as the primary outcome, we can enumerate these regressor permutations exhaustively as follows:

1. Univariate macaque visual brain activity (*M*).
2. Univariate vision model prediction (*V* ).
3. Univariate language model prediction (*L*).
4. Bivariate macaque + vision model (*M* + *V* ).
5. Bivariate macaque + language model (*M* + *L*).
6. Bivariate vision + language model (*L* + *V* ).
7. Trivariate combination of macaque, vision, and language model (*M* + *V* + *L*)

Once (noise-ceiling normalized) scores have been obtained for all 7 of these regressions we can derive the unique and shared (explainable) variances for all 3 of our predictors as follows:

#### Unique variances of *M, V* , *L*

1. *M*_*Unique*_ = (*M* + *V* + *L*) − (*L* + *V* )
2. *V*_*Unique*_ = (*M* + *V* + *L*) − (*M* + *L*)
3. *L*_*Unique*_ = (*M* + *V* + *L*) − (*M* + *V* )

#### 3-way shared variance

4. (*M* + *V* + *L*)_*Shared*_ = *M* + *V* + *L*− 2 * (*M* + *V* + *L*) + *M*_*Unique*_ + *V*_*Unique*_ + *L*_*Unique*_

#### 2-way shared variances

5. (*M* + *V* )_*Shared*_ = *M* + *V* − (*M* + *V* ) − (*M* + *V* + *L*)_*Shared*_
6. (*M* + *L*)_*Shared*_ = *M* + *L*− (*M* + *L*) − (*M* + *V* + *L*)_*Shared*_
7. (*V* + *L*)_*Shared*_ = *V* + *L*− (*V* + *L*) − (*M* + *V* + *L*)_*Shared*_

With or without noise-ceiling normalization, the initial units of these unique and shared variances are proportions of the total variance explained by the trivariate regression (*M* + *V* + *L*). Without noise-ceiling normalization, the units are the proportion of total variance explained. With noise-ceiling normalization, the units are the proportion of total *explainable* variance explained. (Descriptive, natural language versions of these derivations are available in Appendix A2).

## Appendix + Supplementary Information

### A1 Differences in Prediction over Time

In our main analysis of macaque visual activity, we assess for difference between unimodal vision and language models in predicting a single 150ms window of neural activity. This leaves open the question of whether there differences between vision and language model prediction vary across time. Perhaps language model predictions (hypothetically more “conceptual” in nature) peak later than vision model predictions (hypothetically more “perceptual” in nature)?

To answer this, we modeled the time-resolved IT multi unit activity (from 0-500 ms post stimulus onset in overlapping 150 ms bins, sampled every 10ms) in 2 monkeys (Monkey 1 and Monkey 2). Results from this analysis are shown in A1

**Figure A1:**
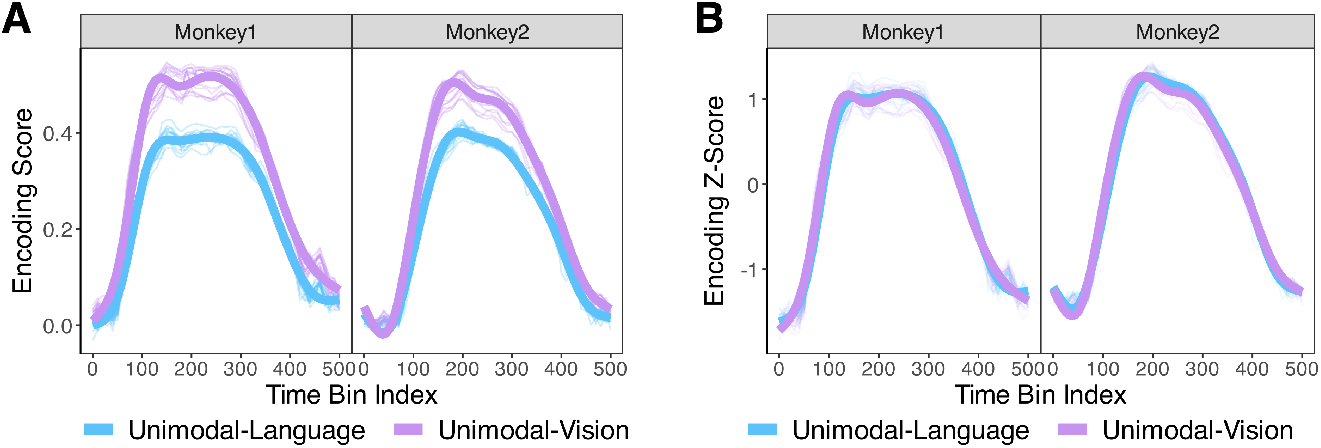
**A** Time-resolved encoding scores for two monkeys from IT multi-unit activity (150 ms bins, 10 ms step size). Thin, semitransparent lines show scores for individual unimodal language (blue) and unimodal vision (purple) models. Thick, opaque lines are the fits of a generalized additive model (GAM) controlling for subject-level variance with an additive term and model-level variance with a random effect term. **B** Time-resolved encoding z-scores obtained after normalizing the scores of each model within their respective modality (allowing for a direct visualization of the overall overlap in the trends across time.))

Results from this analysis make strikingly clear that while the difference between vision and language model prediction is persistent and stable from 100ms to 350ms post stimulus onset, there is *no difference* in the overall trends of each. *Visually*, this parity in overall predictive trend across model modality is most evident in A1B (which depicts the time-varying predictions of each model after their scores are normalized within their respective modality). *Statistically*, this parity is evident in a bootstrapped cross-correlation (lag 0) of *r*=0.995 [.993, .996] in Monkey 1 and .997 [.996, 0.998] in Monkey 2, and the almost identical complexity / effective degrees of freedom (EDF) – 8.95, and 8.96 for vision and language, respectively – in the curves fit by a generalized additive model (thin-plate spline regression) controlling for subject-level variance (with an additive term) and model-level variance (with a random effects term).

In short, while pure language models may predict less variance in macaque visual brain activity than pure visual models *at peak*, the predictions of both modalities seem to increase or decrease in almost-exact parallel across time.

### A2 Further Details on Variance Partitioning

Side-by-side narrative and formulaic descriptions of all terms in the variance partitioning analysis are available in the enumerated lists below. A plot showing side-by-side comparisons of the variance partitioning analysis for both human OTC *and* macaque IT is shown in A2.

#### Unique (unshared) variances

1. *M*_*Unique*_ = (*M* + *V* + *L*) − (*L* + *V* ) The variance in human brain activity explained uniquely by macaque visual brain activity *M* is the trivariate regression (*M* +*V* + *L*), minus the bivariate regression *not* involving the macaque activity: the vision and language models (*L* + *V* ).
2. *V*_*Unique*_ = (*M* + *V* + *L*) − (*M* + *L*) The variance in human brain activity explained uniquely by the vision model is the trivariate regression (*M* +*V* + *L*), minus the bivariate regression *not* involving the vision model: the macaque and language model (*M* + *L*)
3. *L*_*Unique*_ = (*M* + *V* + *L*) − (*M* + *V* ) The variance in human brain activity explained *uniquely* by the language model is the trivariate regression (M + V + L), minus the bivariate regression *not* involving the language model: the macaque and vision model (*M* + *V* ).

#### 3-way shared variance

The variance in human visual brain activty explained *jointly* by all 3 predictors (macaque visual brain activity, vision model, and language model) is the sum of the variance for each individual regression *M* plus *V* plus *L* minus twice the trivariate regression (*M* + *V* + *L*) plus the *unique* variance for each univariate predictor: *M*_*Unique*_ plus *V*_*Unique*_ plus *L*_*Unique*_.

#### 2-way shared variances

4. (*M* + *V* )_*Shared*_ = *M* + *V* − (*M* + *V* ) − (*M* + *V* + *L*)_*Shared*_ The variance in human brain activity explained *jointly* by the macaque visual brain activity and the vision model is the sum of the univariate regressions of each *M* plus *V* minus the bivariate regression of both (*M* +*V* ) minus the variance shared across all (*M* + *V* + *L*)_*Shared*_.
5. (*M* + *L*)_*Shared*_ = *M* + *L*− (*M* + *L*) − (*M* + *V* + *L*)_*Shared*_ The variance in human brain activity explained *jointly* by the macaque visual brain activity and the language model is the sum of the univariate regressions of each *M* plus *V* minus the bivariate regression of both (*M* + *V* ) minus variance shared across all (*M* + *V* + *L*)_*Shared*_.
6. (*V* + *L*)_*Shared*_ = *V* + *L*− (*V* + *L*) − (*M* + *V* + *L*)_*Shared*_ The variance in human brain activity explained *jointly* by the vision and language models is the sum of the univariate regressions of each *V* plus *L* minus the bivariate regression of both (*M* + *V* ) minus the variance shared across all (*M* + *V* + *L*)_*Shared*_

**Figure A2:**
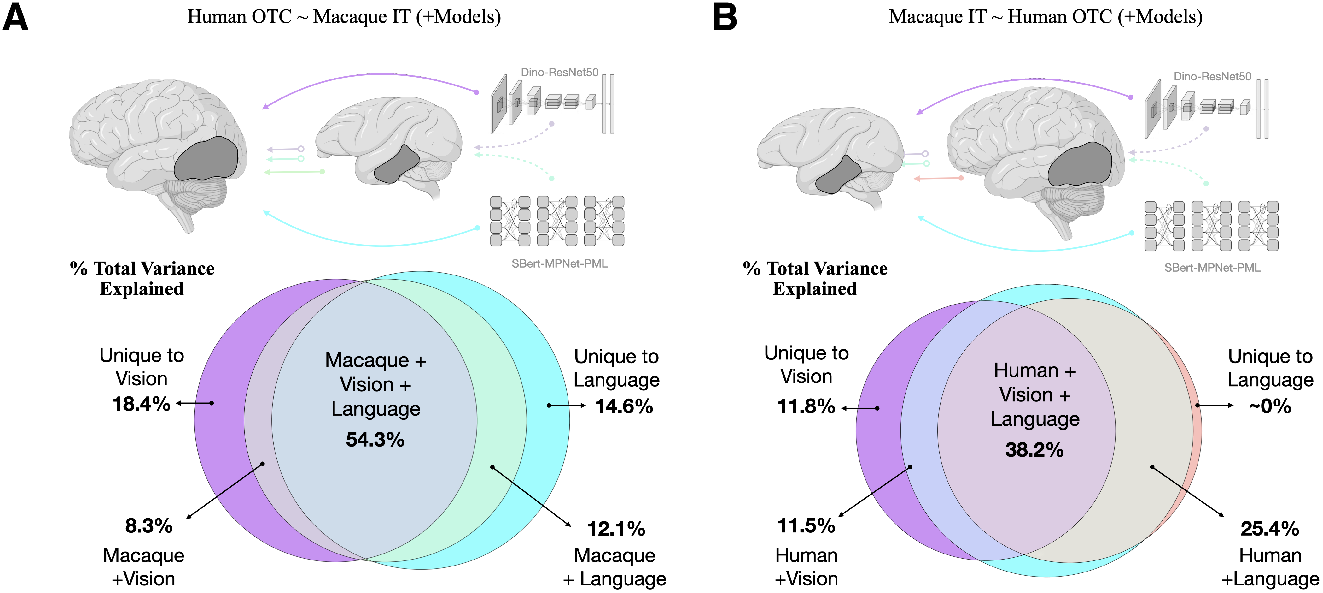
Results from a 3-way variance partitioning analysis designed to quantify the variance in macaque visual brain activity attributable uniquely to vision and uniquely to language, all while accounting for the variance shared with human visual brain activity. **A** is a reproduction (for side-by-side comparison) of the same variance partitioning analysis in 2, without summation of the shared variance terms. **B** is the variance partitioning of macaque IT across the 3 key predictors: human OTC, the most-predictive pure vision-model (Dino-ResNet50) and the most predictive pure-language model (SBERT-MPNet-PML). Apart from the preservation of the individual shared variance terms, all plotting conventions are the same as in Figure 2.

**Figure A3:**
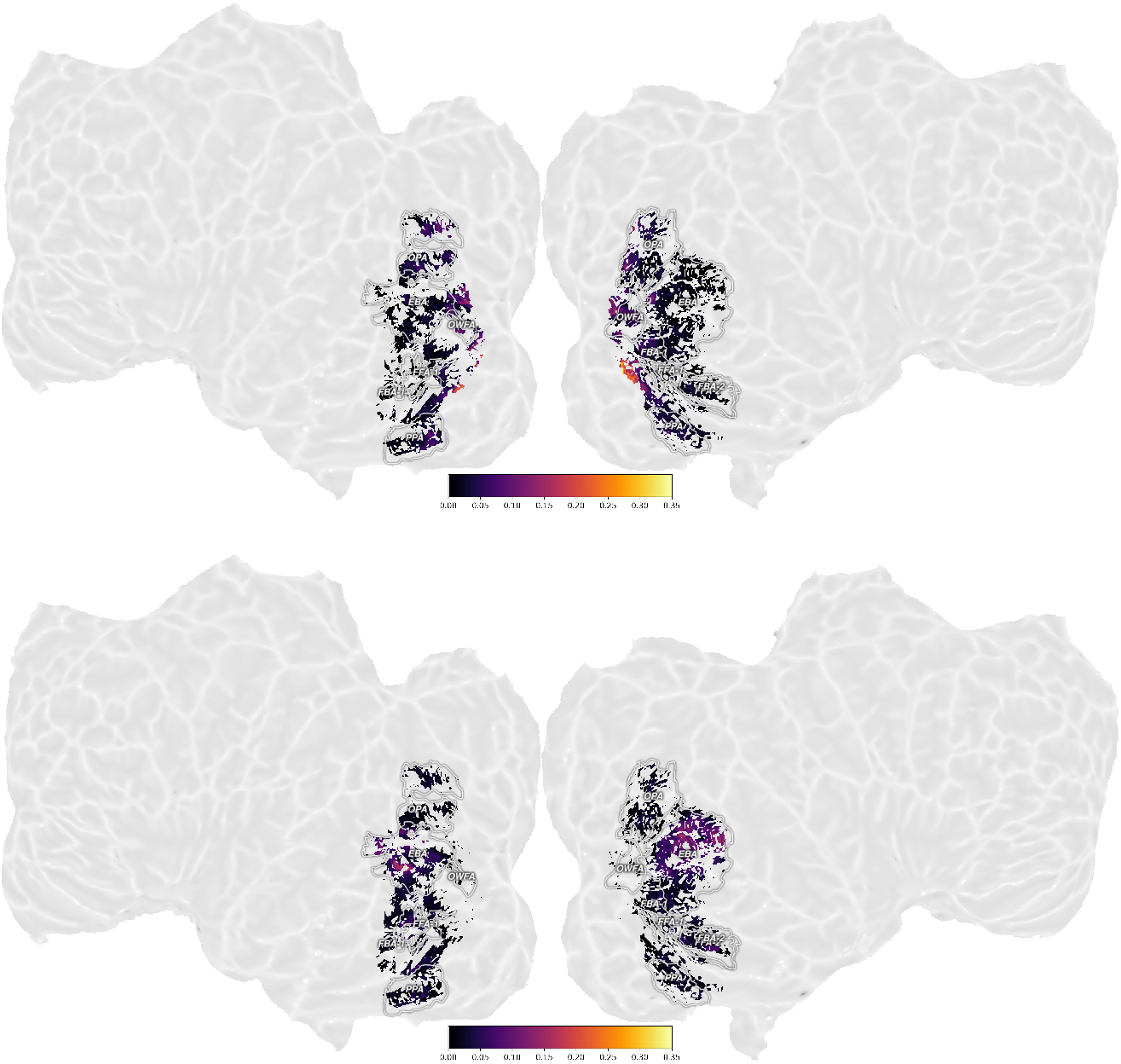
Cortical flat maps showing voxel-wise results of 3-way variance partitioning analysis for Subject 01 (Figures 2, A2A) across 3 key predictors: macaque IT, the most-predictive pure vision-model (Dino-ResNet50) and the most predictive pure-language model (SBERT-MPNet-PML). Top: Voxel-wise unique variance explained by the best vision model. Bottom: Voxel-wise unique variance explained by the best language model.

**Figure A4:**
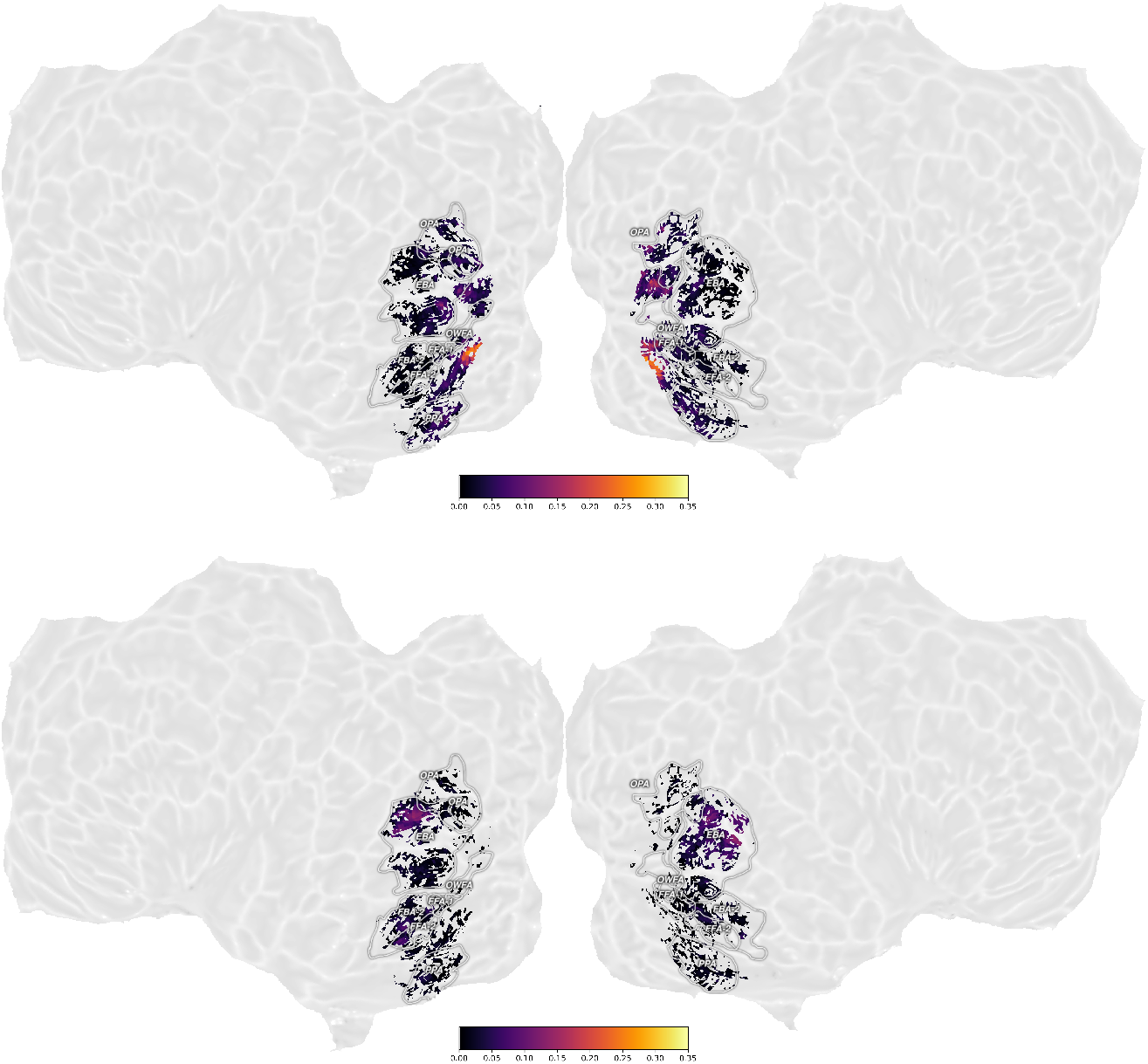
Cortical flat maps showing voxel-wise results of 3-way variance partitioning analysis for Subject 02 (Figures 2, A2A) across 3 key predictors: macaque IT, the most-predictive pure vision-model (Dino-ResNet50) and the most predictive pure-language model (SBERT-MPNet-PML). Top: Voxel-wise unique variance explained by the best vision model. Bottom: Voxel-wise unique variance explained by the best language model.

**Figure A5:**
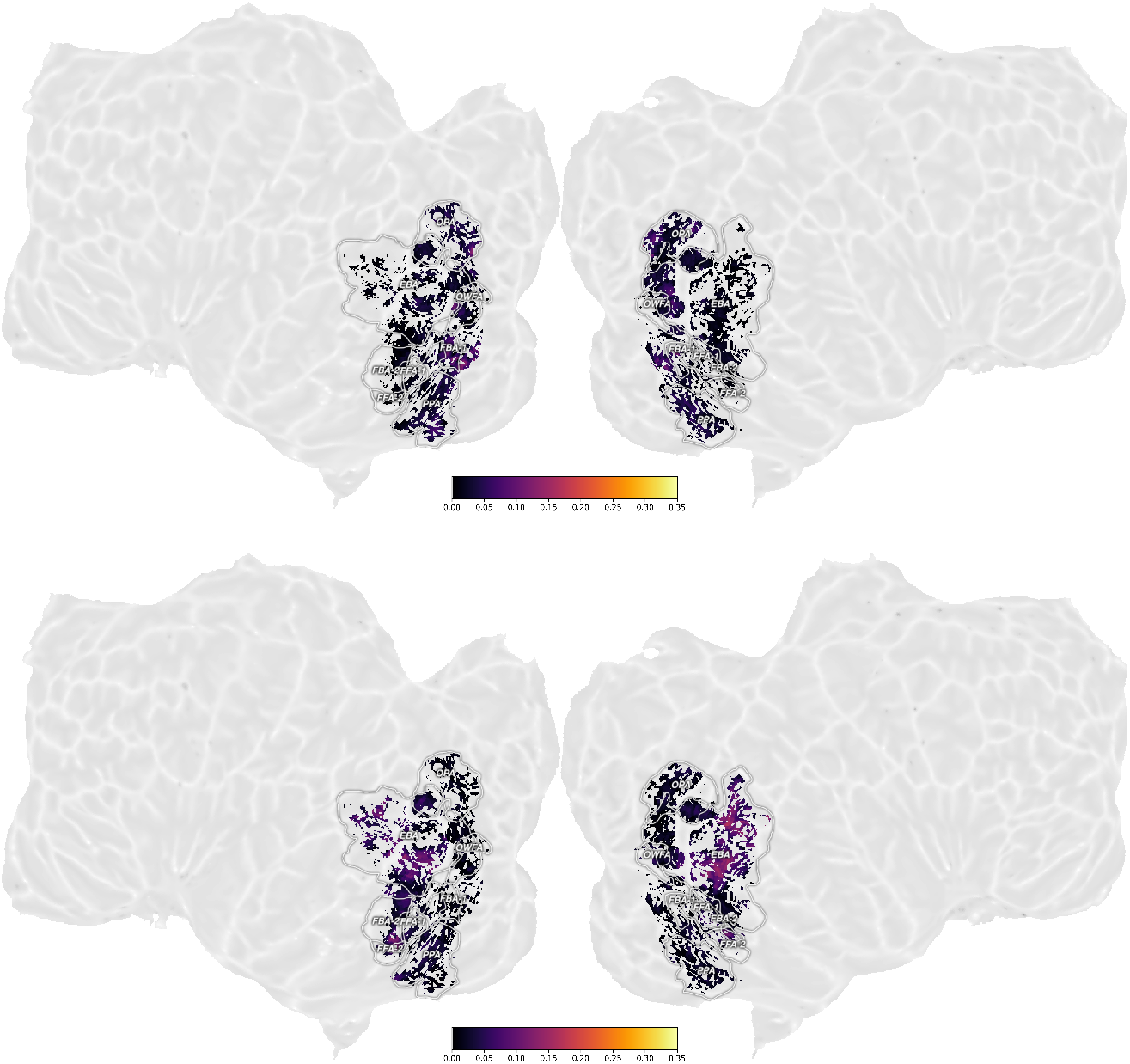
Cortical flat maps showing voxel-wise results of 3-way variance partitioning analysis for Subject 05 (Figures 2, A2A) across 3 key predictors: macaque IT, the most-predictive pure vision-model (Dino-ResNet50) and the most predictive pure-language model (SBERT-MPNet-PML). Top: Voxel-wise unique variance explained by the best vision model. Bottom: Voxel-wise unique variance explained by the best language model.

**Figure A6:**
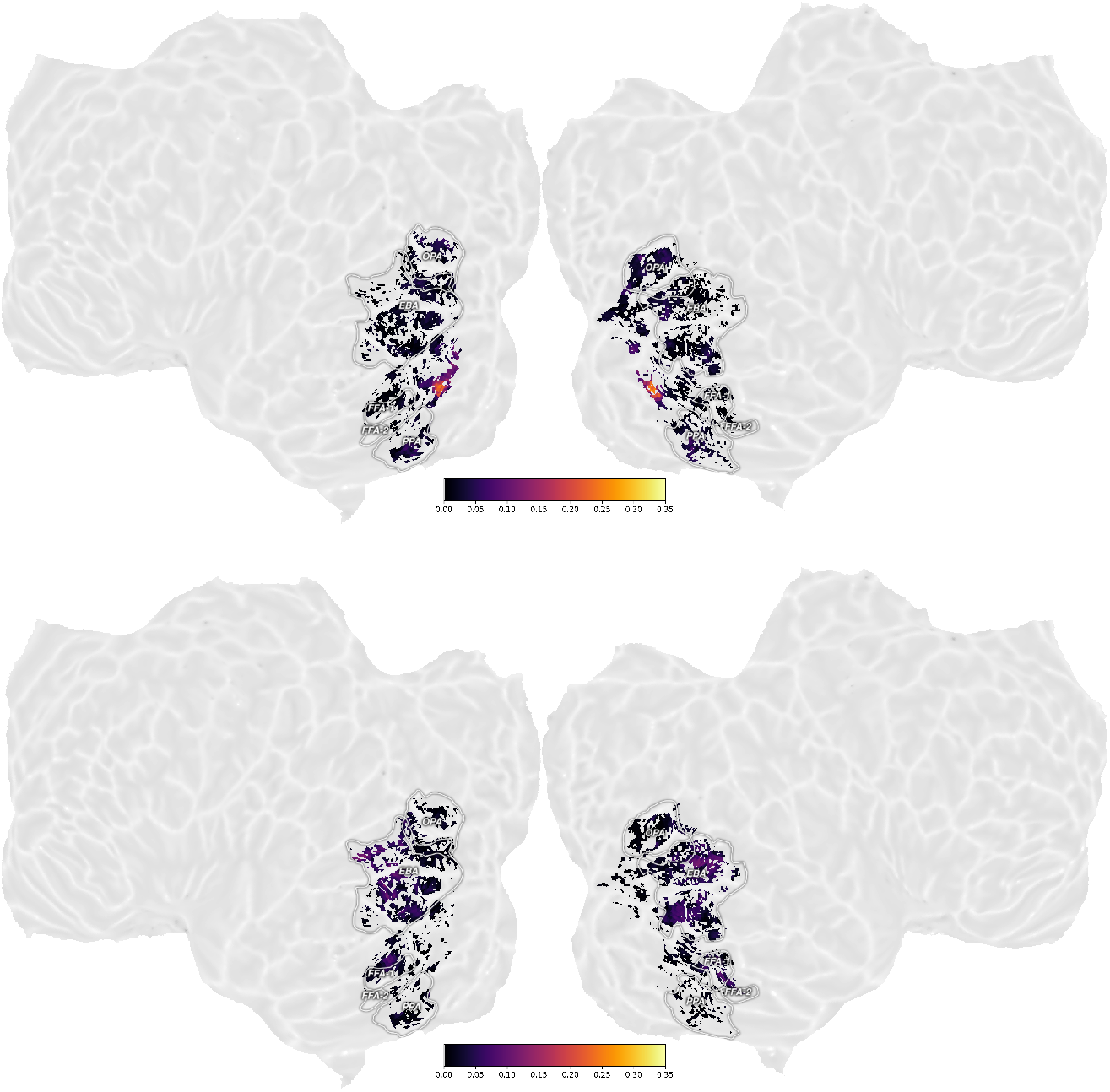
Cortical flat maps showing voxel-wise results of 3-way variance partitioning analysis for Subject 07 (Figures 2, A2A) across 3 key predictors: macaque IT, the most-predictive pure vision-model (Dino-ResNet50) and the most predictive pure-language model (SBERT-MPNet-PML). Top: Voxel-wise unique variance explained by the best vision model. Bottom: Voxel-wise unique variance explained by the best language model.

